# Major improvements to the *Heliconius melpomene* genome assembly used to confirm 10 chromosome fusion events in 6 million years of butterfly evolution

**DOI:** 10.1101/029199

**Authors:** John W. Davey, Mathieu Chouteau, Sarah L. Barker, Luana Maroja, Simon W. Baxter, Fraser Simpson, Mathieu Joron, James Mallet, Kanchon K. Dasmahapatra, Chris D. Jiggins

## Abstract

The *Heliconius* butterflies are a widely studied adaptive radiation of 46 species spread across Central and South America, several of which are known to hybridise in the wild. Here, we present a substantially improved assembly of the *Heliconius melpomene* genome, developed using novel methods that should be applicable to improving other genome assemblies produced using short read sequencing. Firstly, we whole genome sequenced a pedigree to produce a linkage map incorporating 99% of the genome. Secondly, we incorporated haplotype scaffolds extensively to produce a more complete haploid version of the draft genome. Thirdly, we incorporated ~20x coverage of Pacific Biosciences sequencing and scaffolded the haploid genome using an assembly of this long read sequence. These improvements result in a genome of 795 scaffolds, 275 Mb in length, with an L50 of 2.1 Mb, an N50 of 34 and with 99% of the genome placed and 84% anchored on chromosomes. We use the new genome assembly to confirm that the *Heliconius* genome underwent 10 chromosome fusions since the split with its sister genus *Eueides,* over a period of about 6 million years.

## Introduction

Understanding evolution and speciation requires an understanding of genome architecture. Phenotypic variation within a population can be maintained by chromosome inversions (Lowry and Willis 2010; Joron *et al.* 2011; Wang *et al.* 2013) and may lead to species divergence (Noor *et al.* 2001; Feder and Nosil 2009) or to the spread of phenotypes by introgression (Kirkpatrick and Barrett 2015). Genetic divergence and genome composition is affected by variation in recombination rate (Nachman and Payseur 2012; Nam and Ellegren 2012). Gene flow between species can be extensive (Martin *et al.* 2013) and varies considerably across chromosomes (Via and West 2008; Weetman *et al.* 2012).

Describing chromosome inversions, recombination rate variation and gene flow in full requires as close to chromosomal assemblies of the genomes of study species as possible. Recombination rate varies along chromosomes and is influenced by chromosome length (Fledel-Alon *et al.* 2009; Kawakami *et al.* 2014), and inversions are often hundreds of kilobases to megabases long. However, many draft genomes generated with short read technologies contain thousands of scaffolds, often without any chromosomal assignment (Bradnam *et al.* 2013; Michael and VanBuren 2015; Richards and Murali 2015). Where scaffolds are assigned to chromosomes, often a substantial fraction of the genome is left unmapped, and scaffolds are often unordered or unoriented along the chromosomes.

To date, there are 9 published Lepidopteran genomes *(Bombyx mori* (Duan *et al.* 2010), *Danaus plexippus* (Zhan *et al.* 2011), *Heliconius melpomene* (Heliconius Genome Consortium 2012), *Plutella xylostella* (You *et al.* 2013), *Melitaea cinxia* (Ahola *et al.* 2014), *Papilio glaucus* (Cong *et al.* 2015a), *Papilio polytes* and *Papilio xuthus* (both Nishikawa *et al.* 2015), *Lerema accius* (Cong *et al.* 2015b)) and several more available in draft *(Bicyclus anynana, Chilo suppressalis, Manduca sexta, Plodia interpunctella*; see LepBase version 1.0 at http://ensembl.lepbase.org). Of these genomes, only *B. mori, H. melpomene, P. xylostella* and *M. cinxia* have scaffolds with chromosome assignments.

The published *Heliconius melpomene* genome (Heliconius Genome Consortium, 2012; version 1.1 used throughout, referred to as Hmel1.1) contained 4,309 scaffolds (“Hmel1.1”, Figure 1, Table 1), 1,775 of which were assigned to chromosomes based on a linkage map built using 43 RAD-Sequenced F2 offspring (Heliconius Genome Consortium 2012, Supplemental Information S4). The total length of the genome was 273 Mb, including 4 Mb of gaps, with 226 Mb (83%) of the genome assigned to chromosomes. The resulting map has been good enough for many purposes, including estimation of introgression of 40% of the genome between *H. melpomene* and *H. cydno* (Martin *et al.* 2013) and identifying breakpoints between *Heliconius, Melitaea cinxia* and *Bombyx mori* (Heliconius Genome Consortium 2012; Ahola *et al.* 2014). However, for understanding these features and mapping inversions and recombinations, Hmel1.1 has several limitations.

**Figure 1.**
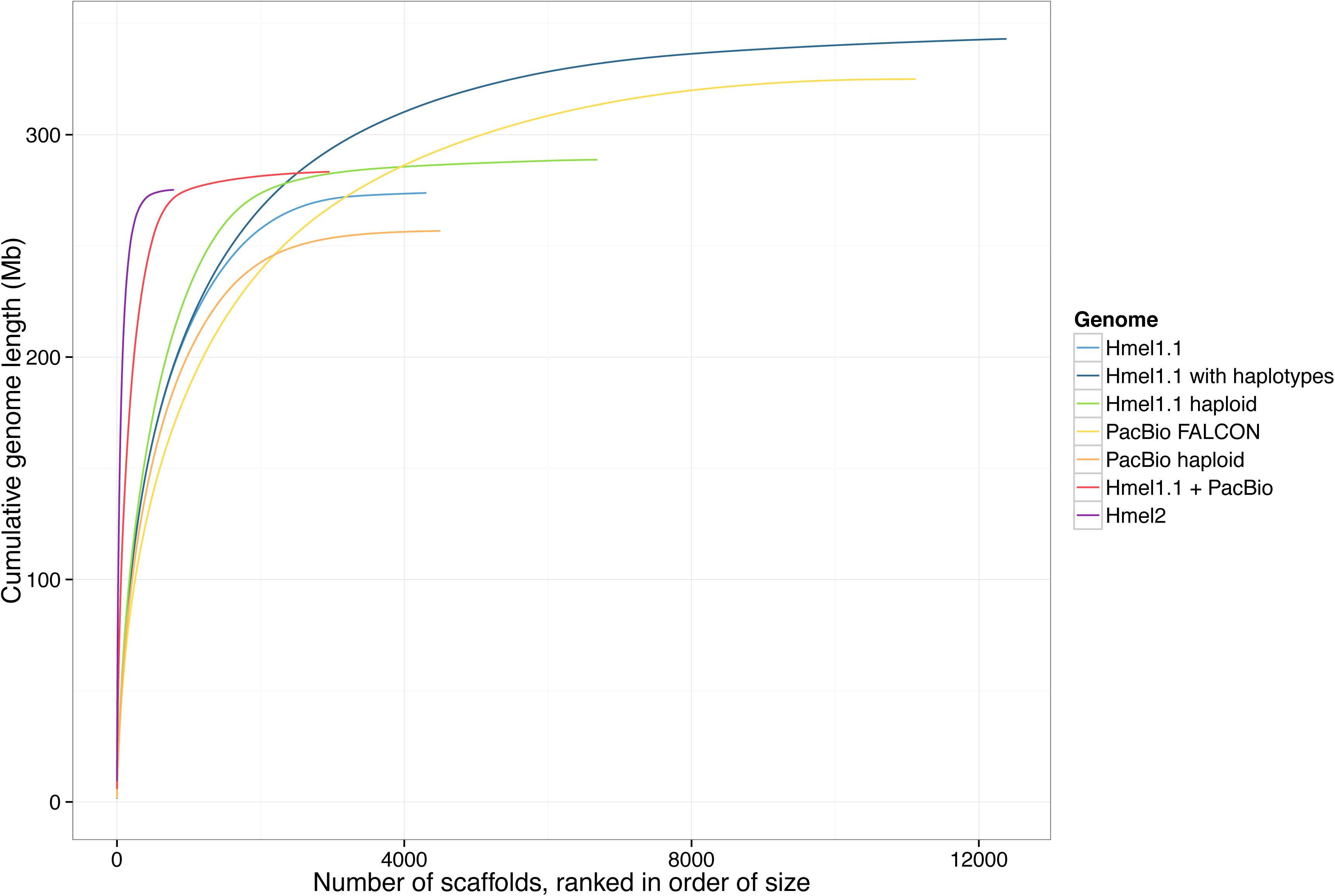
Genome assembly quality. A perfect assembly would appear as an almost straight vertical line. Horizontal plateaus indicate many very small scaffolds. The top right end of each curve shows the number of scaffolds and genome size in the whole assembly. See Table 1 for statistics.

**Table 1.** Statistics for genome assembly versions. Mb, megabases; kb, kilobases; L50, length of scaffold such that 50% of the genome is in scaffolds of this length or longer. N50, number of scaffolds as long as or longer than L50. Colours and names match Figure 1.

| Assembly | Length (Mb) | Scaffolds | Scaffold N50 | Scaffold L50 | Contig L50 |
| --- | --- | --- | --- | --- | --- |
| Hmel1.1 | 273 | 4,309 | 345 | 194 kb | 51 kb |
| Hmel1.1 with haplotypes | 343 | 12,386 | 567 | 128 kb | 33 kb |
| Hmel1.1 haploid | 289 | 6,689 | 346 | 214 kb | 47 kb |
| PacBio FALCON | 325 | 11,121 | 719 | 96 kb | 96 kb |
| PacBio haploid | 256 | 4,565 | 345 | 178 kb | 178 kb |
| Hmel1.1 + PacBio | 283 | 2,961 | 113 | 629 kb | 316 kb |
| Hmel2 | 275 | 795 | 34 | 2.1 Mb | 330 kb |

The original RAD Sequencing linkage map used to place scaffolds on chromosomes in Hmel1.1 was built using the restriction enzyme PstI (cut site CTGCAG), which cuts sites ~10kb apart in the *H. melpomene* genome (32% GC content). Scaffolds shorter than 10kb often did not contain linkage map SNPs and could not be placed on chromosomes. Also, misassemblies could be identified but only corrected to within ~10 kb. With only 43 offspring used in the cross, the average physical distance between recombinations for Hmel1.1 was 320 kb. Scaffolds that could be mapped to a single linkage marker but not more (and so did not span a recombination) could be placed on the linkage map but could not be anchored. Either only one scaffold would be placed at a single marker and could not be oriented, or multiple scaffolds would be placed at a single marker and could not be ordered or oriented. While 226 Mb (83%) of the genome was placed on chromosomes, only 73 Mb (27%) of the genome could be anchored (ordered and oriented). As 17% (46 Mb) of the genome could not be placed on the map, consecutive anchored scaffolds were not joined, as unplaced scaffolds may have been missing in between.

Although the primary Hmel1.1 assembly contained 4,309 scaffolds, an additional 8,077 scaffolds (69 Mb) were identified as haplotypes and removed from the assembly (Heliconius Genome Consortium 2012, Supplemental Information S2.4; “Hmel1.1 with haplotypes”, Figure 1, Table 1). These scaffolds contained 2,480 genes and have been used in several cases to manually bridge primary scaffolds and assemble important regions of the genome (including the Hox cluster, Heliconius Genome Consortium 2012, Supplemental Information S10). It seemed plausible that the assembly would be improved by better genome-wide incorporation of these haplotype scaffolds, rather than their removal.

Since Hmel1.1 was published, long read technologies have matured to the point where high coverage with long reads can be used to produce very high quality assemblies for small or haploid genomes (Berlin *et al.* 2015). Several tools are also available for scaffolding existing genomes with Pacific Biosciences (PacBio) sequence (English et al. 2012; Boetzer and Pirovano 2014). However, these methods are limited by requiring single reads to connect scaffolds, whereas it is likely that many gaps sequenced by PacBio sequencing but missed by Illumina and 454 sequencing (Ross *et al.* 2013) are longer than single reads. An alternative approach is to assemble the PacBio sequence, so that PacBio-unique sequence is retained, and then combine the PacBio assembly with the existing assembly, but tools for doing this have previously been lacking.

Here, we present Hmel2, the second version of the *H. melpomene* genome, which benefits from the use of three techniques to make substantial improvements to the genome assembly: whole genome sequencing of a pedigree, merging of haplotypic sequence, and incorporation of assembled PacBio sequence into the genome.

We have used Hmel2 to test the hypothesis that the *Heliconius* genome underwent 10 chromosome fusions since *Heliconius* split from the neighbouring genus *Eueides* over a period of about 6 million years. It has been known for several decades that all 11 *Eueides* species have 31 chromosomes, whereas *Heliconius* vary from 21 to 56 (Brown *et al.* 1992). It was previously thought that *Heliconius* gradually lost or fused 10 chromosomes via the *Laparus* and *Neruda* genera, which have chromosome numbers between 20 and 31 and had unresolved relationships with *Eueides* and *Heliconius* (Beltrán *et al.* 2007). However, the most recent molecular taxonomy of the Heliconiini (Kozak *et al.* 2015) places *Laparus* and *Neruda* as clades within *Heliconius*, implying that the ancestral chromosome number of *Heliconius* is 21 and suggesting there are no extant species with intermediate chromosome numbers between *Eueides* and *Heliconius*. The change in chromosome number is due to fusions rather than loss, because the 31 chromosomes of *Melitaea cinxia* can be mapped to the 21 chromosomes of *H. melpomene* (Ahola *et al.* 2014). As *Eueides* butterflies also have 31 chromosomes, it seems most likely that these fusions happened since the split between *Eueides* and *Heliconius,* but this has not yet been confirmed. Here, we use a small pedigree of *Eueides isabella* to test whether fusion points between *Eueides* and *Heliconius* match those between *Melitaea* and *Heliconius.*

## Methods

### Preparation of cross

The cross used to build a linkage map for Hmel2 was the same cross used in the original *Heliconius melpomene* genome project (Heliconius Genome Consortium 2012, Supplemental Information section S4). A fourth generation male *H. melpomene melpomene* from an inbred strain was crossed with a female *H. melpomene rosina* (F0 grandmother) from a laboratory strain, both raised in insectaries in Gamboa, Panama. The male was from the same lineage used to produce the Hmel1.1 genome sequence, to ensure the cross was close to the assembly; the female was from a different subspecies to ensure many SNPs were available for use as markers. Two siblings from this F1 were crossed to produce F2 progeny, many of which were frozen at a larval stage. Where possible, sex was determined from wing morphology of individuals that successfully eclosed. Sex of the larval offspring was determined later using sex-linked markers (identified using offspring with known sexes). DNA from the F0 grandmother (the F0 grandfather was lost), two F1 parents and 69 of their F2 offspring was extracted using the DNeasy Blood and Tissue Kit (Qiagen). All samples were prepared as 300 bp insert size Illumina TruSeq libraries except for offspring 11, 16, 17 and 18, which were prepared as Nextera libraries due to low DNA quantities. Libraries were sequenced using 100-bp paired-end reads on an Illumina HiSeq2500 at the FAS Centre for Systems Biology genomics facility, Harvard University. Samples were sequenced over three HiSeq runs. Sequencing failed during sequencing of the second read for two libraries together containing 24 individuals; these libraries were resequenced, but the first run data was still used, with the second read truncated to 65 bases.

### Alignment and SNP calling

Reads for parents and offspring were aligned to Hmel1.1 using Stampy (Lunter and Goodson 2011) version 1.0.23 with options --substitutionrate=0.01 and — gatkcigarworkaround and converted from SAM to BAM format with the SortSam tool from Picard version 1.117 (http://broadinstitute.github.io/picard). Reads were aligned to the primary scaffolds (Hmel1-1_primaryScaffolds.fa) and haplotype scaffolds (Hmel_haplotype_scaffolds.fas) separately. Duplicate reads were removed using the Picard MarkDuplicates tool. Indels were realigned using the RealignerTargetCreator and IndelRealigner tools from GATK version 3.2.2 (DePristo *et al.* 2011). SNPs were called for each individual using the GATK HaplotypeCaller and combined into one final VCF file using GATK GenotypeGVCFs with options --annotateNDA and --max_alternate_alleles 30.

### Conversion of SNPs to marker regions

SNPs were assigned to a marker type according to the calls for the two F1 parents and F0 grandmother (see Table S1 for valid marker types and expected offspring genotypes) or rejected if no valid marker type could be found. SNPs were then rejected if any offspring had an invalid call for the assigned marker type; if the offspring calls failed a root-mean-square test for goodness of fit to expected segregation for the marker type (Perkins *et al.* 2011); if parental genotype quality fell below 99 for heterozygous calls or 60 for homozygous calls; if parental sequencing depth was greater than 85 reads for any parental call; if the SNP had FS (Fisher Strand bias) value greater than 5; or if the SNP had MQ (Mapping Quality) value less than 90. SNP genotypes were converted from GATK format (0/0, 0/1, 1/1) to single letters (A, H, B) for homozygous for allele A (0), heterozygous for alleles A (0) and B (1), and homozygous for allele B (1). Calls were concatenated across all offspring to form a segregation pattern, and phased for each marker type to ensure segregation patterns of each type could be compared.

With millions of remaining markers and low sequencing depth per marker for most offspring, it was impractical to build a linkage map without further reducing the number of SNPs and genotyping errors. Consecutive valid markers of each marker type on each scaffold were therefore converted into consensus markers spanning regions of the scaffold. Scaffolds were split into different regions if more than 25% of offspring differed in their genotype between two consecutive SNPs. For each offspring, the defined scaffold regions were then split into sub-regions by consecutive identical genotype calls, rejecting sub-regions shorter than 100bp (likely due to mis-mapping or poor quality reads). Consensus genotypes were called for each offspring along each sub-region, allowing at most one recombination per offspring per region. At this point, each scaffold features a set of overlapping regions for each valid marker type with consensus genotype calls for each offspring. Valid marker types were grouped into three classes; maternal, where the F1 mother is heterozygous and the F1 father is homozygous; paternal, where the F1 father is heterozygous and the F1 mother is homozygous; and intercross, where both F1 parents are heterozygous (see Table S1 for further details on valid marker types).

### Identification of maternal chromosome prints and paternal markers

As recombination is absent in *Heliconius* females (Turner and Sheppard 1975), a maternal *H. melpomene* linkage map consists of 21 chromosome prints, as all maternal genotypes on the same chromosome are in complete linkage (Jiggins *et al.* 2005). The chromosome prints for Hmel2 were identified by finding scaffold regions with consistent maternal, paternal and intercross markers and then extracting the maternal markers. Markers were labelled consistent when combining maternal and paternal markers and phasing appropriately produced a marker identical to the corresponding intercross marker (because combining markers where only one of each of the parents was heterozygous can result in the pattern produced when both parents are heterozygous). This does not remove all errors, as the same error can occur in multiple marker types. To collapse errors and identify the chromosome prints, log odds (LOD) scores were calculated between each pair of maternal markers and, if a pair of markers had a LOD score below 6, the markers were joined together into one print. 19 of 21 chromosome prints could be identified in this way. By comparing to the set of valid maternal, paternal and intercross markers, scaffold regions with only a valid paternal marker could then be assigned to their corresponding maternal chromosome print; scaffold regions with only a valid intercross marker could be assigned both maternal and paternal markers.

Two chromosomes segregated identically in both F1 parents and so only produced intercross markers, because both parents shared the same variants and so both parents were heterozygous at all loci. These chromosome prints were identified by collapsing intercross markers without matching maternal markers into sets of markers with 6 or fewer different homozygous calls and calculating a consensus of homozygous calls for each set. This produced two sets each with one consensus marker. Paternal markers for regions with one of these markers could then be inferred from the intercross and maternal markers together. This produced a set of 21 maternal chromosome prints and a set of consistent paternal markers with assignments to regions across all scaffolds.

### Linkage map construction

Linkage maps were constructed for each chromosome by ordering paternal markers assigned to each of the 21 maternal chromosome prints iteratively using MSTMap (Wu *et al.* 2008). MSTMap was run with the following options: population_type RIL2, distance_function kosambi, cut_off_p_value 0.000001, no_map_dist 0, no_map_size 0, missing_threshold 1, estimation_before_clustering yes, detect_bad_data yes, objective_function ML.

For each chromosome, an initial map was built using paternal markers each covering more than 200,000 base pairs. If MSTMap returned 2 or more linkage groups, markers were phased to match the first linkage group and the map was built again to produce a single linkage group. Remaining paternal markers were then ordered by the number of base pairs they covered, largest first, and added to the map one by one, rebuilding the map each time. If the new marker was incorporated and introduced a double recombination at that marker in one offspring, that offspring was corrected and the marker was merged into the correct neighbouring marker. If the new marker created a disordered map, or it was added at either end of the map, or it could not be incorporated at all, it was rejected. After all markers had been processed once, further attempts were made to incorporate the rejected markers using the same rules, until an iteration added no new markers to the map.

### *Preprocessing and fixing misassemblies in* Hmel1.1

The primary and haplotype scaffolds of Hmel1.1 were concatenated together and then repeat masked using RepeatMasker 4.0.5 (Smit, AFA, Hubley, R. & Green, P. RepeatMasker 0pen-4.0. 2013-2015 http://www.repeatmasker.org) with the *H. melpomene* version 1.1 repeat library (Hmel.all.named.final.1-31.lib, Lavoie *et al.* 2013) as input and with options -xsmall and -no_is. Candidate misassemblies in Hmel1.1 were identified by detecting discontinuities in linkage map markers across genomic scaffolds, and then manually validated to identify the smallest possible breakpoint based on marker SNPs, including SNPs that were rejected from linkage map construction but could be assigned to one of the two markers around the breakpoint. Long misassembled regions (~5kb or greater) were retained as separate scaffolds but most misassembled regions were discarded. Breakpoints that spanned two contigs or contained an entire contig were likely due to scaffolding errors; in these cases the scaffold was broken at the gap. If an entire contig was contained within a breakpoint, with no additional SNP to link it to the markers on either side, it was discarded.

Misassemblies corrected in version 1.1 were also revisited (Heliconius Genome Consortium 2012, Supplementary Information S4.6). The linkage map used to place scaffolds for version 1.1 was built using RAD Sequencing data, with samples cut with the PstI restriction enzyme. This produces sites roughly 10 kilobases apart, which meant that many breakpoints were not identified accurately. Each of the misassemblies was reconsidered here, with all of the previously broken scaffolds remerged and new breakpoints defined based on the whole genome mapping data.

Errors in the linkage map were identified during the merging and reassembly processes described below. A list of linkage map errors was constructed and erroneous blocks removed and corrected using a script, clean_errors.py.

### Merging genome

HaploMerger version 20120810 (Huang *et al.* 2012) was used to collapse haplotypes in the *H. melpomene* genome. A scoring matrix for LASTZ (as used within HaploMerger) was generated using the lastz_D_Wrapper.pl script with --identity=94. This scoring matrix was used for all runs of HaploMerger, including for the PacBio genome (see below). HaploMerger was run with default settings except for setting --size=20 in all_lastz.ctl, targetSize=5000000 and querySize=400000000 in hm.batchA.initiation_and_all_lastz, and haploMergingMode=“updated” in hm.batchF.refine_haplomerger_connections_and_Ngap_fillings.

Several scripts were written to make running HaploMerger easier. The new script runhm.pl executes one iteration of HaploMerger, running batch scripts A, B, C, E, F and G, renaming output scaffolds with a given prefix, producing a final FASTA file concatenating merged scaffolds and unmerged scaffolds, and generating summary statistics (using summarizeAssembly.py in PBSuite 14.9.9, http://sourceforge.net/projects/pb-jelly/, English *et al.* 2012) and an AGP file for the final FASTA (using bespoke script agp_from_fasta.py). The HaploMerger script hm.batchG.refine_unpaired_sequences was not used for the initial Hmel1.1 and PacBio assembly merges, retaining all potentially redundant scaffolds in case they could be used for scaffolding later, but it was used to merge the haploid Hmel1.1 assembly with the haploid PacBio assembly. The new script batchhm.pl runs runhm.pl iteratively until HaploMerger fails to merge any further scaffolds. It also runs a set of additional new scripts map_merge.py, transfer_merge.py and transfer_features.py, that document where the original genome parts are in the new genome. The map_merge.py script takes HaploMerger output and documents where the input genome scaffolds are in the merged output genome. The transfer_merge.py script takes this transfer information and another transfer file, for example between the original version 1.1 *H. melpomene* genome and the input genome, and computes the transfer from the original genome to the output genome. The transfer_features.py script then transfers linkage map markers, genes and misassembly information to the new genome.

HaploMerger sometimes merges scaffolds incorrectly, but has several mechanisms for users to manually edit its output. The hm.nodes file, which contains detected overlaps between scaffolds, can be manually annotated, with incorrect merges marked to be rejected. The revised hm.nodes file is then passed through the batchE script to update the merged scaffolds to ignore the incorrect merges. Incorrect merges in the *Heliconius* genome could be detected by comparing against the linkage map data. A list of scaffolds that should not be merged was constructed over multiple merge attempts and runhm.pl was used to edit the hm.nodes and run the batchE script automatically.

HaploMerger merges scaffolds based on overlaps and reports the parts contributing to merged scaffolds in the hm.new_scaffolds file, including which of the two overlapping parts has been included in the new genome. These choices sometimes broke genes, whereas choosing the other part would retain the annotated gene. runhm.pl can also take a GFF file as input and check for broken genes in hm.nodes and hm.new_scaffolds, rejecting nodes if they break manually curated genes, and swapping parts in an overlap if it prevents gene breakage. It then runs the batchE and batchF to update the merged scaffolds. The Hmel1.1 GFF files (heliconius_melpomene_v1.1_primaryScaffs_wGeneSymDesc.gff3 and Hmel1-0_HaplotypeScaffolds.gff) were concatenated and passed to runhm.pl to avoid as many breakages of Hmel1.1 genes as possible.

### Pacific Biosciences sequencing, error correction and assembly

A pupa from the *H. melpomene* genome strain from Gamboa, Panama was dissected and DNA extracted using the QIAGEN HMW MagAttract kit. This pupa was taken after four generations of inbreeding, and came from the same generation as the F0 father used to construct the pedigree reported here, and the generation before the individuals used for the genome sequence itself. A Pacific Biosciences (PacBio) SMRTbell 25kb needle sheared library was constructed, size selected with 0.375x SPRI beads and sequenced using P4/C2 chemistry (180 minute movie).

PacBio subreads were self-corrected with PBcR (in Celera assembly v8.3, Berlin *et al.* 2015), run with options -length 200, -genomeSize 292000000) and separately corrected with the original genome strain Illumina (Sequence Read Archive accession SRX124669), 454 shotgun (SRX124544) and 454 3kb mate-pair (SRX124545) sequencing data (using option -genomeSize 292000000). Self-corrected and genome-strain-corrected reads were concatenated into one read set and assembled with FALCON (https://github.com/PacificBiosciences/falcon, commit bb63f20d500efa77f930c373105edb5fbe37d74b, 2 April 2015) with options input_type=preads, length_cutoff=500, length_cutoff_pr=500, pa_HPCdaligner_option=“-v -dal4 -t16 -e.70 -l1000 -s1000”, ovlp_HPCdaligner_option=“-v -dal32 -t32 -h60 -e.95 -l500 -s1000, pa_DBsplit_option=“-x500 -s50”, ovlp_DBsplit_option=“-x500 -s50”, falcon_sense_option=“--output_multi --min_idt 0.70 -- min_cov 4 --local_match_count_threshold 2 --max_n_read 100 --n_core 6”, overlap_filtering_setting=“--max_diff 40 --max_cov 60 --min_cov 2 --bestn 10”.

The FALCON assembly was merged iteratively to exhaustion using batchhm.pl as with version 1.1 of the *H. melpomene* genome (see previous section). Misassemblies in the PacBio assembly were identified using the same methods as Hmel1.1 and the merge was repeated several times to remove these misassemblies.

### Scaffolding and gap filling with PacBio assembly

The final, ‘haploid’ merged Hmel1.1 and PacBio genomes were merged together using runhm.pl. For this final merge, gap filling in hm.batchF.refine_haplomerger_connections_and_Ngap_fillings was turned on, and runhm.pl edited hm.new_scaffolds to always select portions from the Hmel1.1 genome over portions from the PacBio genome, to preserve as much of the Hmel1.1 genome as possible and use the PacBio genome for scaffolding only. Also, hm.batchG.refine_unpaired_sequences was run and the refined FASTA output used, to remove as many redundant sequences from the resulting merged genome as possible. Finally, runhm.pl was run on the merged Hmel1.1+PacBio genome, to generate a set of nodes for use in scaffolding. Linkage map markers and genes were transferred to this final merged genome with transfer_features.py.

### Cleaning merged assembly and ordering scaffolds along chromosomes

The Hmel1.1+PacBio merged genome was cleaned and ordered with reference to the linkage map markers. Scaffolds coming from the PacBio assembly alone were removed. If HaploMerger incorporates some portion P of a scaffold S into a merged scaffold, it retains the remaining portions of the scaffold as new scaffolds. These remaining portions were labelled offcuts. Offcuts were removed from the genome if they contained no markers on the linkage map, or if they mapped to the same chromosomal location as the merged scaffold containing their original portion P, assuming that the offcut is part of a haplotype. However, some offcuts that mapped to different chromosomal locations were retained, as they were often long portions of scaffolds that had been misassembled. Scaffolds were also removed if they mapped to a marker that mapped within a larger scaffold that featured surrounding markers; for example, if scaffold A has markers 1,2,3, and scaffold B has marker 2 only, scaffold B was removed as an assumed haplotype.

Scaffolds were ordered along chromosomes based on their linkage markers. Pools of scaffolds were defined containing one or more scaffold. If a pool contained a single scaffold that bridged multiple consecutive markers, the scaffold could be ordered and oriented and so was labelled ‘anchored’. A pool containing a single scaffold spanning only a single marker could be ordered on the chromosome but not oriented, and so was labelled ‘unoriented’. A pool containing multiple scaffolds at a single marker was labelled ‘unordered’, as the scaffolds could be neither ordered or oriented against each other.

This order was refined by using the nodes (overlaps between pairs of scaffolds) identified by HaploMerger in the merged Hmel1.1+PacBio genome. HaploMerger does not use all the nodes it identifies, relying on a scoring threshold to reject low-affinity overlaps. While this is sensible when merging over a whole-genome, many of these nodes proved to be useful when considering single pools or neighbouring pools of scaffolds. Scaffolds that had a connecting node in a scaffold in a neighbouring pool that would mean that the scaffold was completely contained in the neighbouring scaffold were removed as likely haplotypes, providing that candidate haplotype scaffolds longer than 10kb had a %alignment greater than 50% and candidate haplotype scaffolds shorter than 10kb had %alignment greater than 25%. If neighbouring scaffolds had an overlapping node at their ends, or were bridged via nodes to a PacBio scaffold, they were ordered and oriented next to each other in the genome, connecting the scaffolds with a 100bp gap.

Consecutive anchored scaffolds were connected together into one scaffold. This was not done during scaffolding for Hmel1.1, as with only 86% of the genome scaffolded it was assumed that large scaffolds may have been missing between anchored scaffolds. However, with 98% of the genome mapped for version 2, it was felt the connection of anchored scaffolds with a gap was reasonable.

After each chromosome was assembled, a set of unmapped scaffolds remained. These scaffolds were retained if they had a maternally informative marker but no paternally informative marker (and so could be placed on the chromosome but not ordered on it), or if they featured a gene. Otherwise, they were removed from the final genome.

### Annotation transfer

Using transfer_features.py (see above), the Hmel1.1 gene annotation could be transferred directly to Hmel2. However, this revealed a number of avoidable gene breakages, where a haplotype scaffold had been incorporated in place of a primary scaffold, but the sequence was still the same or similar. CrossMap (version 0.1.8, http://crossmap.sourceforge.net) was used to transfer as many remaining annotations by alignment as possible, using HaploMerger to produce a chain map of Hmel1.1 against Hmel2 to use as input to CrossMap.

### Identifying Eueides and Melitaea chromosome fusion points

*Eueides isabella* subspecies (male *dissoluta,* female *eva)* were crossed in insectaries in Tarapoto, Peru. Parents were whole genome sequenced and 21 F1 offspring were RAD sequenced using the PstI restriction enzyme on an Illumina HiSeq 2500. Offspring were separated by barcode using process_radtags from version 1.30 of Stacks (Catchen et al. 2011). Parents and offspring were aligned to Hmel2 using the same alignment pipeline described above except using GATK version 3.4-0 and Picard tools version 1.135. UnifiedGenotyper was used for SNP calling rather than HaplotypeCaller as HaplotypeCaller does not perform well with RAD sequencing data. SNPs where the father was homozygous, the mother was heterozygous (or, for the Z chromosome, had a different allele to the father) and the offspring all had genotypes were identified. The resulting segregation patterns were sorted by number of SNPs. The most common segregation patterns and mirrors of these patterns were identified as chromosome prints, as no other patterns appeared at large numbers of SNPs, except for where all offspring were homozygous, or where the patterns were genotyping errors from the chromosome prints. The positions of the SNPs for each chromosome print were then examined to identify fusion points, with clear transitions from one segregation pattern to another visible for all ten fused chromosomes.

The fusion points in *Heliconius* relative to *Melitaea cinxia* were identified by running HaploMerger on a merge of Hmel2 and the *M. cinxia* version 1 genome superscaffolds (Melitaea_cinxia_superscaffolds_v1.fsa.gz, downloaded from http://www.helsinki.fi/science/metapop/research/mcgenome2downloads.html on 14 July 2015). Overlaps (nodes) detected by HaploMerger between Hmel2 scaffolds and *M. cinxia* superscaffolds were used to confirm synteny based on known chromosomal assignments of *M. cinxia* superscaffolds. All fusion points could be identified using this method except for *Heliconius* chromosome 20, which was confirmed using progressiveMauve (as used by Ahola *et al.* (2014) to confirm synteny between *H. melpomene, M. cinxia and B. mori;* Mauve version 2.4.0 Linux snapshot 2015-02-13 used, Darling *et al.* 2010).

### Lepidopteran genome statistics

Lepidopteran genomes compared in Table 2 were downloaded from LepBase v1.0 (http://ensembl.lepbase.org) on October 2, 2015, except for *Danaus plexippus* version 3 (http://monarchbase.umassmed.edu/download/Dpgenomev3.fasta.gz), *Papilio polytes* (http://papilio.nig.ac.jp/data/Ppolytesgenome.fa.gz) and *Papilio xuthus* (http://papilio.nig.ac.jp/data/Pxuthusgenome.fa.gz). Summary statistics were calculated using summarizeAssembly.py in PBSuite 14.9.9 (http://sourceforge.net/projects/pb-jelly/, English *et al.* 2012) and bespoke script genome_kb_plot.pl, used to calculate N50s and make plots for Figure 1 and Figure S3. BUSCO values were calculated using BUSCO v1.1b1 with the set of 2675 arthropod genes (Simão *et al.* 2015).

**Table 2.**
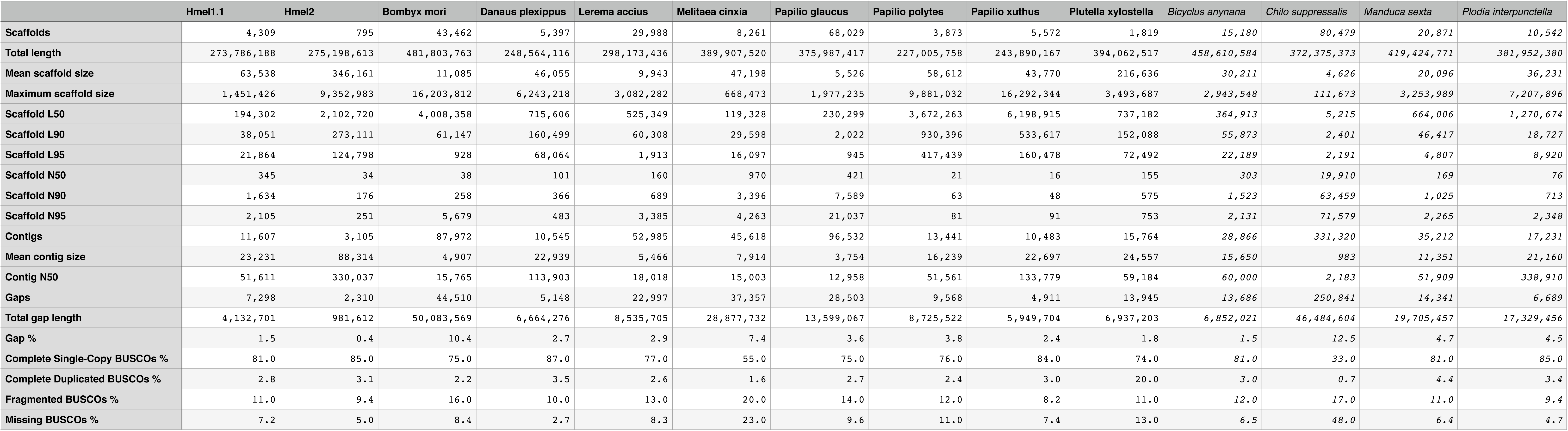
Genome assembly statistics for Hmel1.1, Hmel2 and other published and unpublished Lepidopteran genomes. See Table 1 for definitions of L50 and N50. BUSCO (Benchmarking Universal Single-Copy Ortholog) values are based on a set of 2675 arthropod BUSCOs (Simão *et al.* 2015). Complete Duplicated BUSCOs are included in the count of Complete Single-Copy BUSCOs. See Methods for details of genomes and calculation of statistics. Statistics in italics are for draft, unpublished genomes and should not be taken as representative of the final genomes when they are published.

### Data availability

The Hmel2 genome is available from LepBase v1.0 (http://ensembl.lepbase.org). A distribution containing the genome and many supplementary files will be made available from butterflygenome.org but if not available at time of review can be found on Dropbox at https://www.dropbox.com/sh/ke92gsnts5gbp5g/AAAoWOJTgBP6Sxu7ElPBQusa?dl=0. Sequence reads from the *H. melpomene* and *E. isabella* crosses are available from European Nucleotide Archive (ENA) accession PRJEB11288. Pacific Biosciences data is available from ENA accession ERP005954. All bespoke code is available on GitHub at https://github.com/johnomics/Heliconius_melpomene_version_2. A Dryad repository containing the Hmel2 distribution, a frozen version of the GitHub repository, VCF files for the *H. melpomene* and *E. isabella* crosses, marker databases, and intermediate genome versions for Hmel1-1 and the PacBio assemblies will be made available on acceptance of the manuscript, but can currently be found on Dropbox at https://www.dropbox.com/sh/l4xp1r920zjuuvm/AAAQq9cI46HKfDrIiP3lhtmma?dl=0.

## Results

### Whole genome sequence genetic map

A genetic map of a full-sib cross between *H. melpomene melpomene* x *H. melpomene rosina* was constructed to place scaffolds from version Hmel1.1 of the *H. melpomene* genome on to chromosomes. The F0 grandmother, F1 parents and 69 offspring were whole genome sequenced and aligned to Hmel1.1 (Table S2). 17.2 million raw SNPs were filtered down to 2.9 million SNPs and converted into 919 unique markers (Table S1). The linkage map built from these markers has 21 linkage groups and a total map length of 1,364.23 cM (Figure 2). 2,749 of 4,309 primary scaffolds and 4,062 of 8,077 haplotype scaffolds contained marker SNPs, adding up to 268 Mb (98%) of the primary sequence and 57 Mb (83%) of the haplotype sequence.

**Figure 2.**
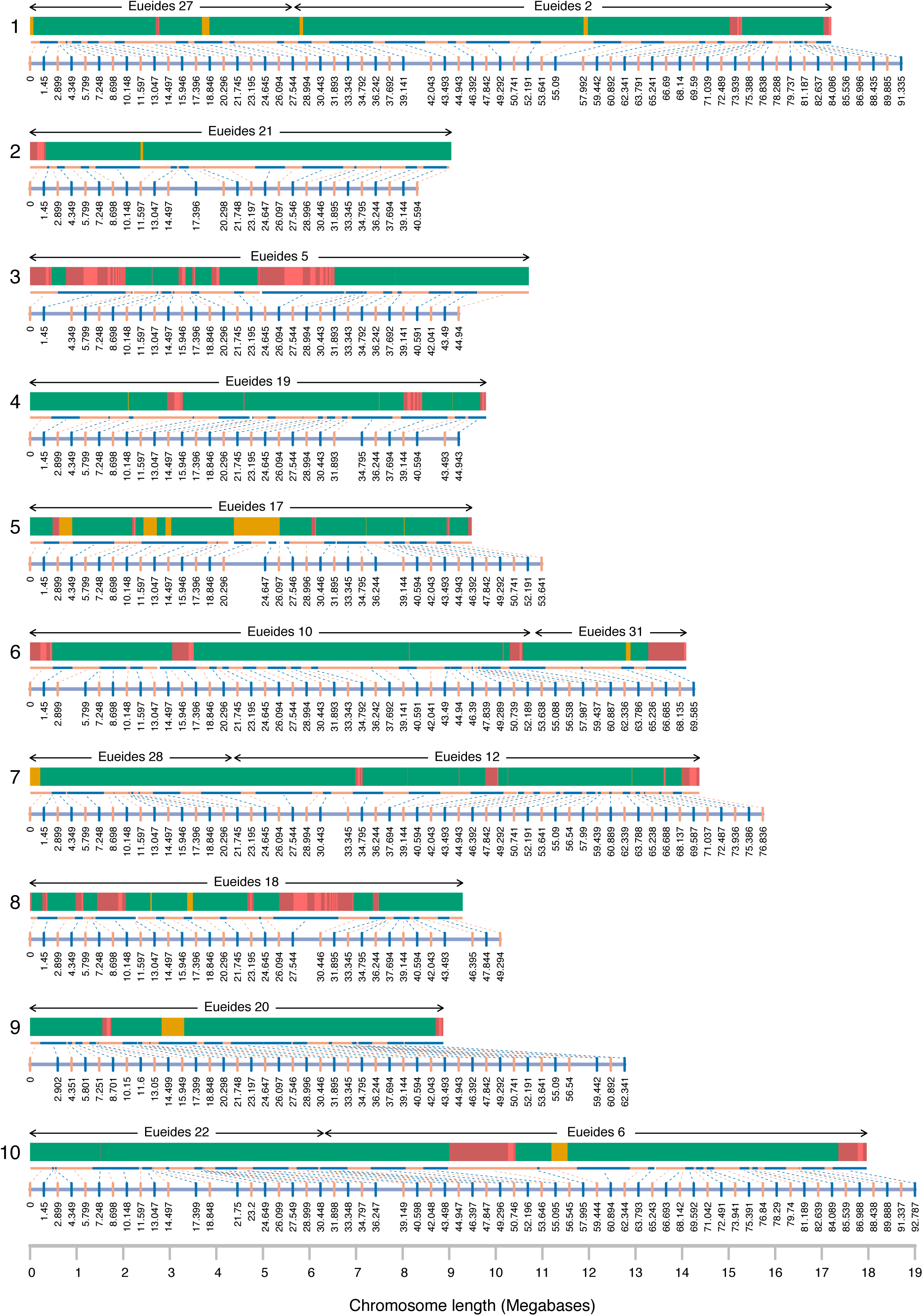
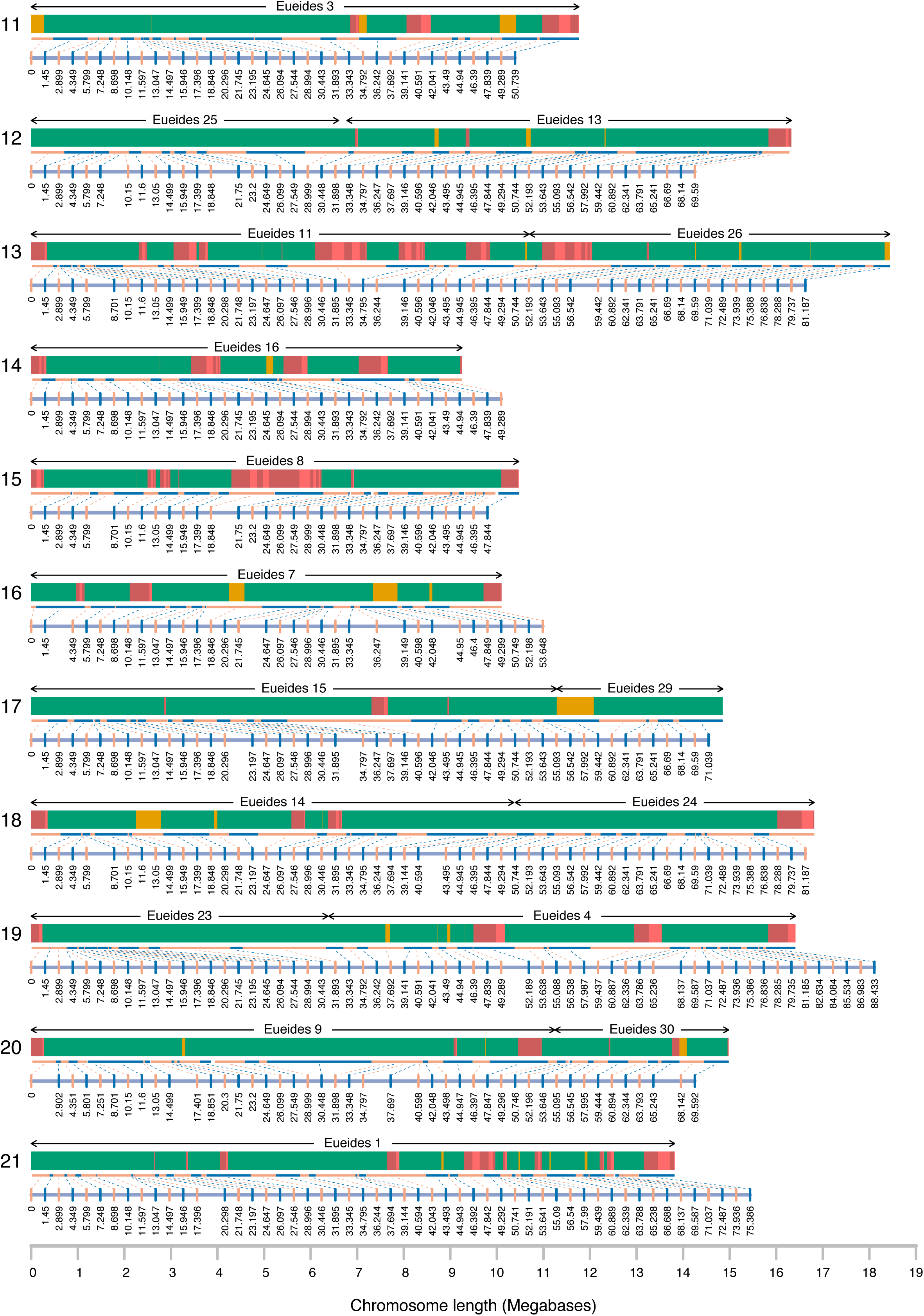
The Hmel2 genome assembly. Chromosome numbers shown on left. Each chromosome has a genetic map and a physical map. Linkage markers (alternating blue and orange vertical lines) connect to physical ranges for each marker (alternating blue and orange horizontal lines) scaled to maximum chromosome length (x-axis at the bottom of each page). Scaffolds are shown in green (anchored), orange (one unoriented scaffold placed at a marker) and alternating light and dark red (multiple unordered scaffolds placed at one marker). Red scaffolds at each marker are arbitrarily ordered by length. *Eueides* chromosome synteny is shown above each chromosome (see Figure 4).

In addition to mapping the majority of the genome sequence to chromosomes, whole genome sequencing of a pedigree allows very accurate detection of crossovers and misassemblies. Identical SNPs could be concatenated into linkage blocks across scaffolds. For example, across the scaffold containing the B/D locus, which controls red patterning in *Heliconius* (Baxter *et al.* 2008; Reed *et al.* 2011), 6 crossovers were called with an average gap of 344 bp between linkage blocks; a misassembly at the end of the scaffold was called with a gap of 2.9 kb (Figure 3). Across the genome, crossover and misassembly gaps have a mean size of 2.2 kb (SD 3.7 kb), all unmapped regions (crossover and misassembly gaps, unmapped scaffold ends or whole unmapped scaffolds) have mean size 2.5 kb (SD 5.1 kb), whereas mapped regions have mean size 28.4 kb (SD 62.7 kb) (see Figure S1 for distributions).

**Figure 3.**
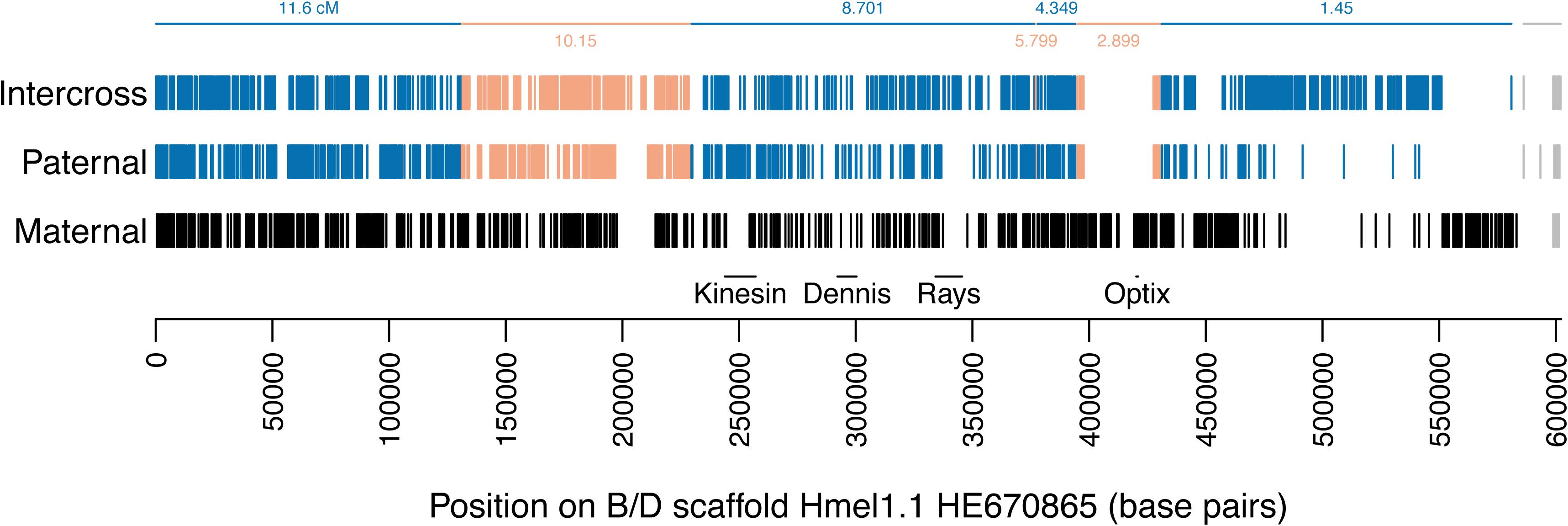
SNPs across the B/D locus scaffold for the major marker types Maternal (F1 mother heterozygous, F1 father homozygous), Paternal (F1 father heterozygous, F1 mother homozygous) and Intercross (both F1 parents heterozygous); see Table S1 for marker type details. Kinesin, Dennis, Rays and Optix are major features of the locus (Baxter *et al.* 2008, Reed *et al.* 2011). Vertical lines, SNPs; horizontal lines, linkage map marker ranges (cf Figure 2). SNP colours: black, maternal pattern for chromosome 18; alternating blue and orange, linkage map markers from 1.45 cM to 11.6 cM on chromosome 18 (cf Figure 2); grey; misassembly, now on chromosome 16.

Based on this linkage information, 380 misassemblies were corrected in the genome. This included revisiting the 149 misassemblies fixed for Hmel1.1 (Heliconius Genome Consortium 2012, Supplementary Information S4.6) to more accurately identify the breakpoints for these misassemblies, and fixing 231 new misassemblies.

### Haplotype merging and scaffolding with PacBio sequencing

The Hmel1.1 primary and haplotype scaffolds were merged together using HaploMerger, iterating 9 times until no further scaffolds could be merged, avoiding gene breakages where possible and reverting merges where they conflicted with the linkage map. This produced a haploid genome containing 6,689 scaffolds, length 289 Mb, L50 214 kb (“Hmel1.1 haploid”, Figure 1, Table 1).

23x coverage of the *H. melpomene* genome was generated using PacBio sequencing. These sequence reads were error-corrected once using the original Illumina and 454 data from the genome and again using self-correction (Table S3). The two error-corrected read sets were combined and assembled together using FALCON to produce an initial assembly of 11,121 scaffolds with L50 96 kb and total length 325 Mb (“PacBio FALCON”, Figure 1, Table 1).

The initial PacBio assembly was merged to itself iteratively using HaploMerger to produce a haploid PacBio assembly (“PacBio haploid”, Figure 1, Table 1). The haploid Hmel1.1 genome and haploid PacBio genome were then merged using HaploMerger to scaffold the two genomes together. This final merge was checked against the linkage map and 470 misassemblies in the original PacBio assembly were fixed, requiring the two PacBio merging steps to be repeated several times. The final haploid PacBio genome had 4,565 scaffolds, L50 178 kb, total length 256 Mb; the Hmel1.1+PacBio merged assembly had 2,961 scaffolds, L50 629 kb, total length 283 Mb (Figure 1, Table 1).

### Ordering of scaffolds on chromosomes

Linkage information was transferred to the Hmel1.1+PacBio merged assembly and used to place the resulting scaffolds on chromosomes, anchoring scaffolds wherever possible, connecting consecutive anchored scaffolds, and removing remaining haplotypic scaffolds (see Methods for details). Further scaffolds were joined by searching for connections to PacBio scaffolds unused by HaploMerger during the merge process. This left 641 scaffolds (274 Mb) placed on chromosomes (98.7% of the genome), with a further 869 scaffolds (3.6 Mb) unplaced. 154 (1.1 Mb) of the unplaced scaffolds were retained as they contained genes or had chromosome assignments (but no placement within the chromosome), and the remaining 715 scaffolds (2.5 Mb, 0.9%) were discarded.

The final genome assembly, Hmel2, has 795 scaffolds, length 275.2 Mb, L50 2.1 Mb (Figure 1, Table 1, Figure 2, Table 2), with 231 Mb (84%) anchored and 274 Mb (99%) placed on chromosomes (Figure S2). This compares well with the other published Lepidopteran genome assemblies to date (Table 2, Figure S3).

### Improved assembly of major loci

The assembly of major adaptive loci is greatly improved in Hmel2, with all scaffolds containing known adaptive loci substantially extended and most gaps filled. The yellow colour pattern locus Yb, previously on a 1.33 Mb scaffold, is now on a 1.96 Mb scaffold; the red pattern BD locus scaffold has increased from 602 kb to 1.89 Mb and is now gap-free; the K locus, previously spread over two scaffolds totalling 173 kb, is now on a single 3 Mb scaffold; the Ac locus, previously on three scaffolds totalling 838 kb is now on a single 7.4 Mb scaffold; and the Hox cluster, previously manually assembled into 7 scaffolds covering 1.4 Mb (Heliconius Genome Consortium 2012, Supplementary Information S10), is now a single scaffold covering 1.3 Mb, with some misassembled material reassigned elsewhere. Full details of major locus locations in Hmel1.1 and Hmel2 (based on loci from Nadeau *et al.* 2014) can be found in Table S4, with three previously unmapped minor loci now placed on chromosomes.

### *Chromosome fusions between* Eueides *and* Heliconius

To identify chromosome fusion points between *Eueides* and *Heliconius,* chromosome prints for the 31 *Eueides* chromosomes were discovered using RAD Sequencing data from an *E. isabella* cross aligned to the Hmel2 genome (Table S5). Synteny between *Heliconius* and *Eueides* is clear on all chromosomes, with 11 unfused and 10 fused *Heliconius* chromosomes (Figure 4). The *Eueides* fusion points all fall within the *Melitaea* fusion points reported by Ahola et al. (2014) and confirmed against Hmel2 here (Table S6), indicating that these fusions occurred since the split between *Eueides* and *Heliconius.* Major colour pattern loci and other adaptive loci (Nadeau et al. 2014) are not near to fusion points, with the exception of the *H. erato* locus Ro, which is 73 kb away from the chromosome 13 fusion point (Figure 4, Table S4).

**Figure 4.**
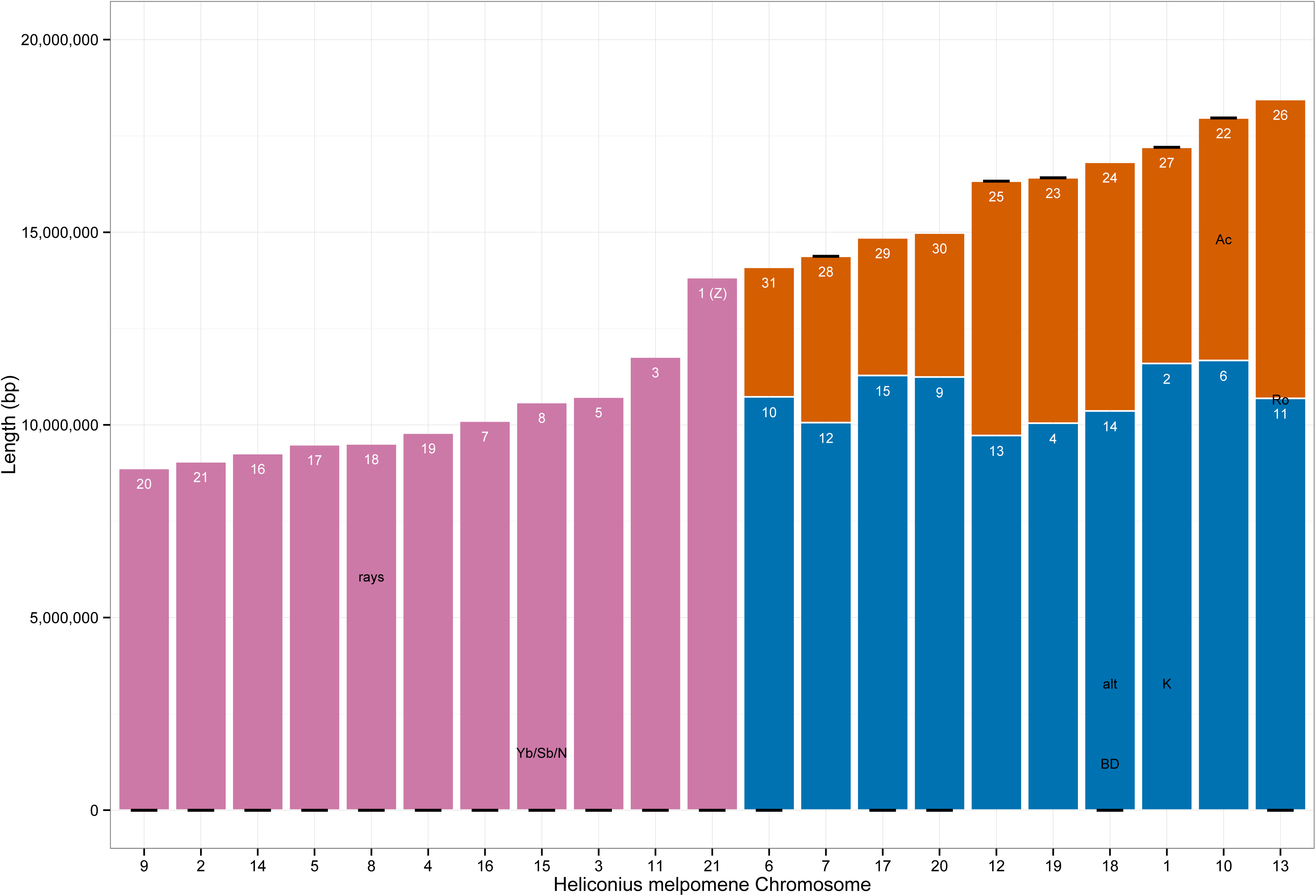
Chromosome fusions in *H. melpomene.* Chromosomes of *H. melpomene* ordered by length. Unfused *Heliconius* chromosomes in pink; fused *Eueides/Melitaea* chromosomes in orange and blue, longest chromosome of each pair in blue. *Melitaea* chromosome numbers in white. Black line, beginning of *H. melpomene* chromosome in Hmel2. Black labels, loci known to be associated with colour pattern features or altitude (alt) in *H. melpomene* or *H. erato* (Nadeau *et al.* 2014).

As noted by Ahola et al. (2014), the shorter *Melitaea* chromosomes (22-31) are all involved in fusions. The longer *Melitaea* autosome in each fusion pair in *Heliconius (Melitaea* 2, 4, 6, 9-15; mean length 10.7 Mb, SD 688 kb) does not, on average, differ substantially in length to unfused autosomes *(Melitaea* 3, 5, 7, 8, 16-21; mean length 9.9 Mb, SD 894 kb). In contrast, the shorter *Melitaea* autosomes in each fusion pair in *Heliconius (Melitaea* 22-31) have mean length 5.4 Mb (SD 1.5 Mb), suggesting a bimodal distribution with the long *Melitaea* autosomes, both fused and unfused, clustering together into one group and the short fused *Melitaea* autosomes clustering into a second group.

## Discussion

### Genome assembly improvements

Many long range technologies are now available for improvement of existing draft genomes. Deep coverage with long reads can be sufficient for producing almost complete *de novo* assemblies (Berlin *et al.* 2015) and additional technologies such as optical mapping can substantially improve genome scaffolding and identify complex structural variants (Pendleton *et al.* 2015, English *et al.* 2015). However, it remains unclear how well these technologies will work with highly heterozygous non-model organisms.

Here, we show that even a small amount of PacBio data (~20x coverage) was sufficient to substantially improve the *H. melpomene* genome. Indeed, the assembly of the PacBio data alone was comparable in quality to our initial draft assembly constructed with Illumina, 454 and mate pair sequencing (Heliconius Genome Consortium 2012; compare lines “Hmel1.1 with haplotypes” and “PacBio FALCON” in Table 1 and Figure 1). We expect that increasing this coverage could have produced a very high quality genome with no additional data.

However, this does not deal with heterozygosity across the genome and the resulting generation of many haplotypic scaffolds, a problem for most species and particularly for insects (Richards *et al.* 2015). As sequencing methods improve and true haplotypes can be assembled, it is hoped that full diploid genomes can be produced, and several efforts are already moving towards this (Church et al. 2015; https://github.com/ekg/vg). We hope that in the near future it will be possible to assemble a diploid reference graph for *H. melpomene,* perhaps with the haplotypes reported here. However, as we wanted to preserve contiguity with Hmel1.1, which was already a composite of both haplotypes, Hmel2 remains a composite haploid genome.

HaploMerger has proved to be a very versatile assembly tool. In addition to having many options for varying the merging process and for manually accepting or rejecting merges, HaploMerger is almost unique among similar tools in reporting where it has placed parts of the original genome in the new genome. This has allowed us to write scripts to transfer linkage map information and genes to new genome versions directly and automatically, without having to map the original genome scaffolds to the new genome separately and possibly erroneously (although we have used this approach to map genes that couldn’t be transferred directly). We could then accept or reject merges where they introduced misassemblies that conflicted with the linkage map or broke genes, and iterate the use of HaploMerger to collapse as many scaffolds as possible. This allowed us to use HaploMerger to scaffold the existing *Heliconius* genome with our novel PacBio genome, by treating the two ‘haploid’ genomes as two haplotypes in one diploid genome. We could then modify the HaploMerger output to prefer the original Hmel1.1 genome over the PacBio genome, only using the PacBio genome for scaffolding, and so preserve our original assembly and annotation wherever possible.

Hmel2 is not complete; it does not contain a W chromosome, and no chromosome is assembled into a single scaffold. The incomplete assemblies may be partially due to errors in haplotype merging. The detailed linkage mapping information available for most scaffolds increases our confidence that primary and haplotype scaffolds have been accurately placed, but it may be that merging haplotypes has collapsed or removed repetitive material. The final genome size of 275 Mb is lower than the flow cytometry estimate of 292 Mb +/− 2.4 Mb (Jiggins et al. 2005). Remaining gaps between scaffolds and failures to order scaffolds may be due to incorrect assembly of haplotypes at the ends of scaffolds, or due to genuine incompatibilities between the many individual butterflies that have contributed to the genome sequence, making it impossible to find overlaps or connections between these ends. Several hundred small scaffolds remain in the genome, which are likely to be misassemblies of repetitive elements, but no clear metric could be found that excluded or integrated these scaffolds. However, as the positions of removed haplotypes have been recorded, it may be possible to reintegrate this material with further analysis of particular regions of the genome. Further manual inspection of existing data, PCRs across scaffold ends, additional long read sequencing, or additional cross sequencing or optical mapping will hopefully resolve many of these remaining assembly problems.

### Is Heliconius speciation rate driven by chromosome fusions?

Chromosome number varies widely in the Lepidoptera (Robinson 1971) and gradual transitions from one number to another occur frequently. Lepidopteran chromosomes are believed to be holocentric (Wolf *et al.* 1994), which may make it easier for chromosome fusions and fissions to spread throughout a population (Melters *et al.* 2012). However, the fusion of 20 chromosomes into 10 over 6 million years is the largest shift in chromosome number in such a short period across the Lepidoptera (Ahola et al. 2014, Figure 3A). Also, given the supposed ease of chromosome number transitions, it is unusual that chromosome number in the Nymphalinae and Heliconiinae is stable at 31 chromosomes for the majority of species, in contrast to all other subfamilies where chromosome number tends to fluctuate gradually and widely (Ahola et al. 2014, Figure 3B). While *Heliconius* species do vary in chromosome number, the majority still have 21 chromosomes, with substantial variations only found in derived clades (Brown et al. 1992; Kozak et al. 2015). It is not just the transition in chromosome number but also the stability of chromosome number before and after the transition that requires explanation.

The difference in chromosome number confirmed here is a major difference between the *Heliconius* and *Eueides* genera which may make these genera an excellent system for studying macroevolution and speciation. Kozak et al. (2015) demonstrated that speciation rate in *Heliconius* is significantly higher than in *Eueides,* but the rate in both genera is more or less stable and does not obviously relate to geological events or adaptive traits. The difference in chromosome number may contribute to explaining this difference in speciation rate, and might provide a null hypothesis for comparison with potential adaptive explanations based on colour pattern, host plant preference, geographic ranges and other traits.

Restriction of recombination facilitates speciation in the presence of gene flow (Butlin 2005). One of the major mechanisms for restricting recombination are chromosome inversions, where opposing alleles can become linked together and then become fixed in different populations (Kirkpatrick *et al.* 2006; Farré et al. 2013; Kirkpatrick *et al.* 2015). However, other methods of restricting recombination may produce similar effects.

Recombination rate is negatively correlated with chromosome length, although the relationship is complex (Fledel-Alon *et al.* 2009; Kawakami *et al.* 2014). In many species, one obligate crossover is required for successful meiosis, inflating recombination rate in short chromosomes. However, beyond a certain length, recombination rate increases roughly linearly with chromosome length (Kawakami *et al.* 2014). It is unclear whether these relationships will hold in Lepidoptera, which may have no obligate crossovers, as females do not recombine and meiosis requires the formation of a synaptonemal complex rather than recombination (Wolf *et al.* 2014).

It is possible that recombination rate along fused chromosomes in *Heliconius* has decreased considerably compared to their shorter, unfused counterparts in *Eueides* (and *Melitaea),* particularly on the shorter chromosomes. This may have enabled linked pairs of divergently selected loci to accrue more easily in *Heliconius* than in *Eueides,* making the process of speciation more likely (Nachman and Payseur 2012, Brandvain *et al.* 2014). This hypothesis could be tested by generating population sequence for *Eueides* species to compare to existing *Heliconius* population data (such as Martin *et al.* 2013), and by modelling speciation rates in the face of different recombination rates. Such a model could predict speciation rate differences between the genera, but full testing would also require the generation of accurate recombination rates in both genera. The system is particularly well suited for testing speciation rate effects because the set of 10 unfused autosomes can act as a control; the hypothesis predicts that recombination rate will not have changed substantially on these chromosomes.

This hypothesis demonstrates the pressing need to generate full, chromosomal genomes for *Eueides* and other *Heliconius* species; genome size in *H. erato* is ~393 Mb (Tobler et al 2005), very similar to *M. cinxia,* but roughly 100 Mb larger than *H. melpomene.* Unpublished draft genome sequences of *Eueides tales* and other *Heliconius* species suggest genome sizes similar to *H. erato* or larger, with *H. melpomene* being one of the smallest *Heliconius* genomes (data not shown). Measuring recombination rate for other species against the *H. melpomene* genome alone is therefore unlikely to be accurate and may not allow for accurate model fitting. However, with additional genomes in hand, we believe these genera may provide a useful test case for the influence of genome architecture on speciation and molecular evolution.

## Acknowledgements

JWD is funded by a Herchel Smith Postdoctoral Research Fellowship. PacBio sequencing was carried out by Paul Coupland and Richard Durbin at the Sanger Institute, supported by the ERC grant numbers 339873, the Wellcome Trust grant number 098051 and JWD’s Herchel Smith funding. *H. melpomene* cross sequencing was carried out at the Harvard FAS Center for Systems Biology core facility and funded by BBSRC grant number G006903/1. *E. isabella* cross sequencing was carried out by Sylviane Moss at the Gurdon Institute. We thank Jenny Barna for computing support. Alignment and SNP calling was performed using the Darwin Supercomputer of the University of Cambridge High Performance Computing Service (http://www.hpc.cam.ac.uk/), provided by Dell Inc. using Strategic Research Infrastructure Funding from the Higher Education Funding Council for England and funding from the Science and Technology Facilities Council.

## Author contributions

JWD and CJ conceived the study. JWD designed the analyses, wrote the software, extracted DNA for PacBio sequencing, and wrote the paper. FS, KKD and JM extracted DNA, prepared libraries and whole genome sequenced the pedigree. LM and SWB bred the *H. melpomene* cross. MC and MJ bred the *Eueides isabella* cross. SLB extracted DNA and made RAD libraries for the *E. isabella* cross. All authors read and commented on the manuscript.

---

**Figure S1.**
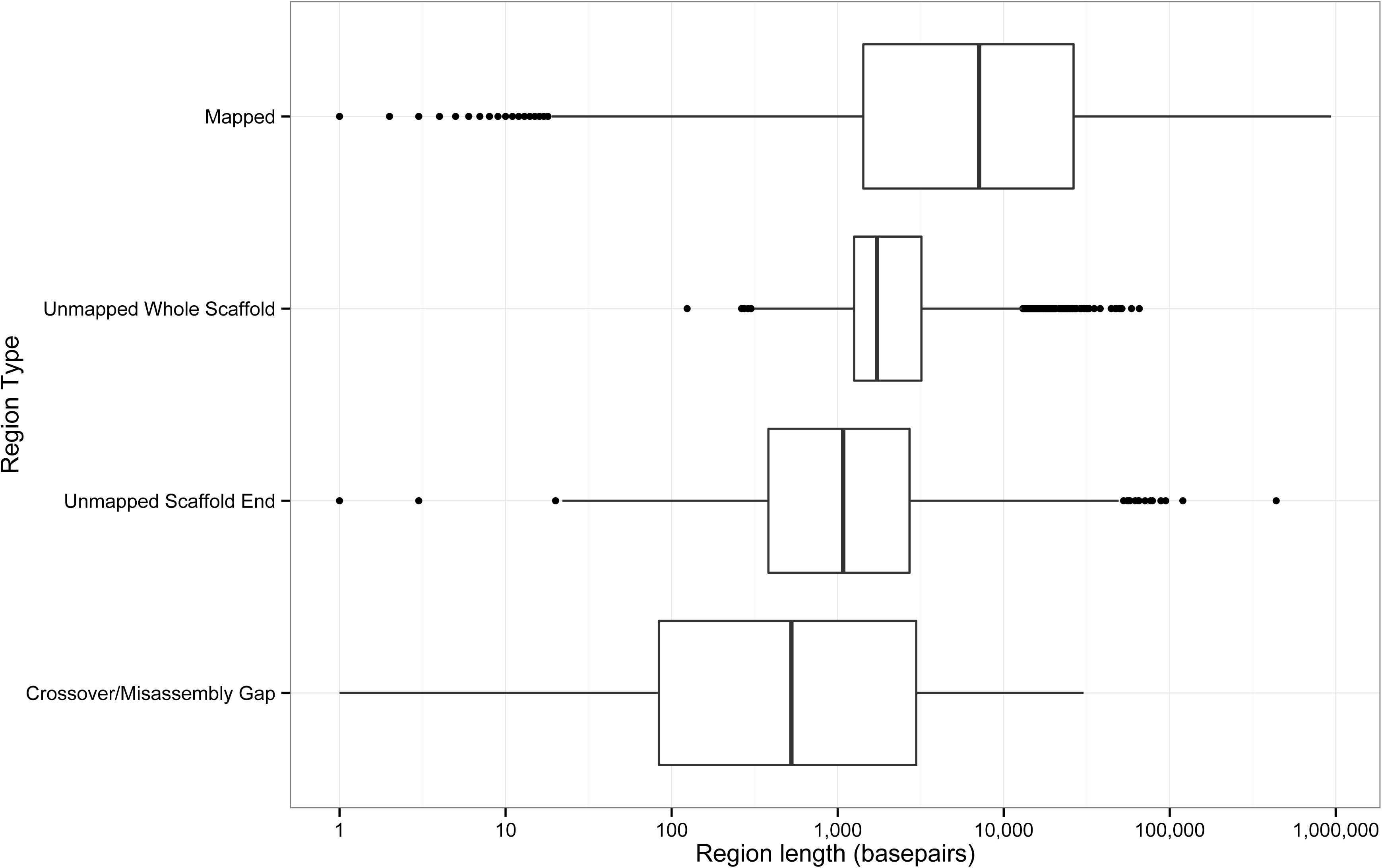
Ranges of mapped and unmapped region lengths across all Hmel1.1 scaffolds. Crossover/Misassembly Gaps occur within scaffolds between markers, either consecutive on one chromosome (Crossover) or distant on one chromosome or on different chromosomes (Misassembly).

**Figure S2.**
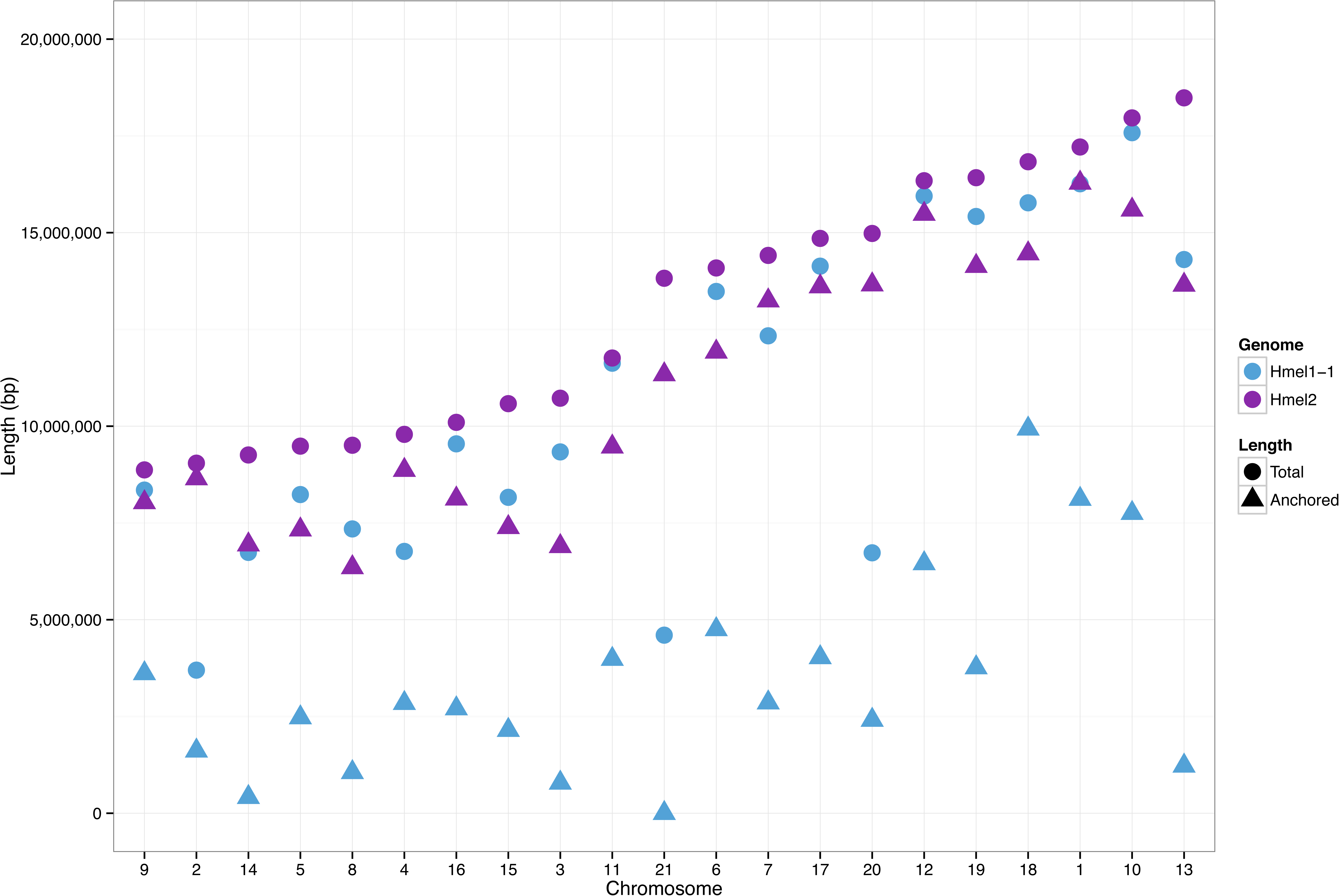
Length of genome assembly placed on chromosomes (Total) and anchored (ordered and oriented, green in Figure 2) on chromosomes (Anchored), for Hmel1.1 and Hmel2. Chromosomes ordered by total length in Hmel2.

**Figure S3.**
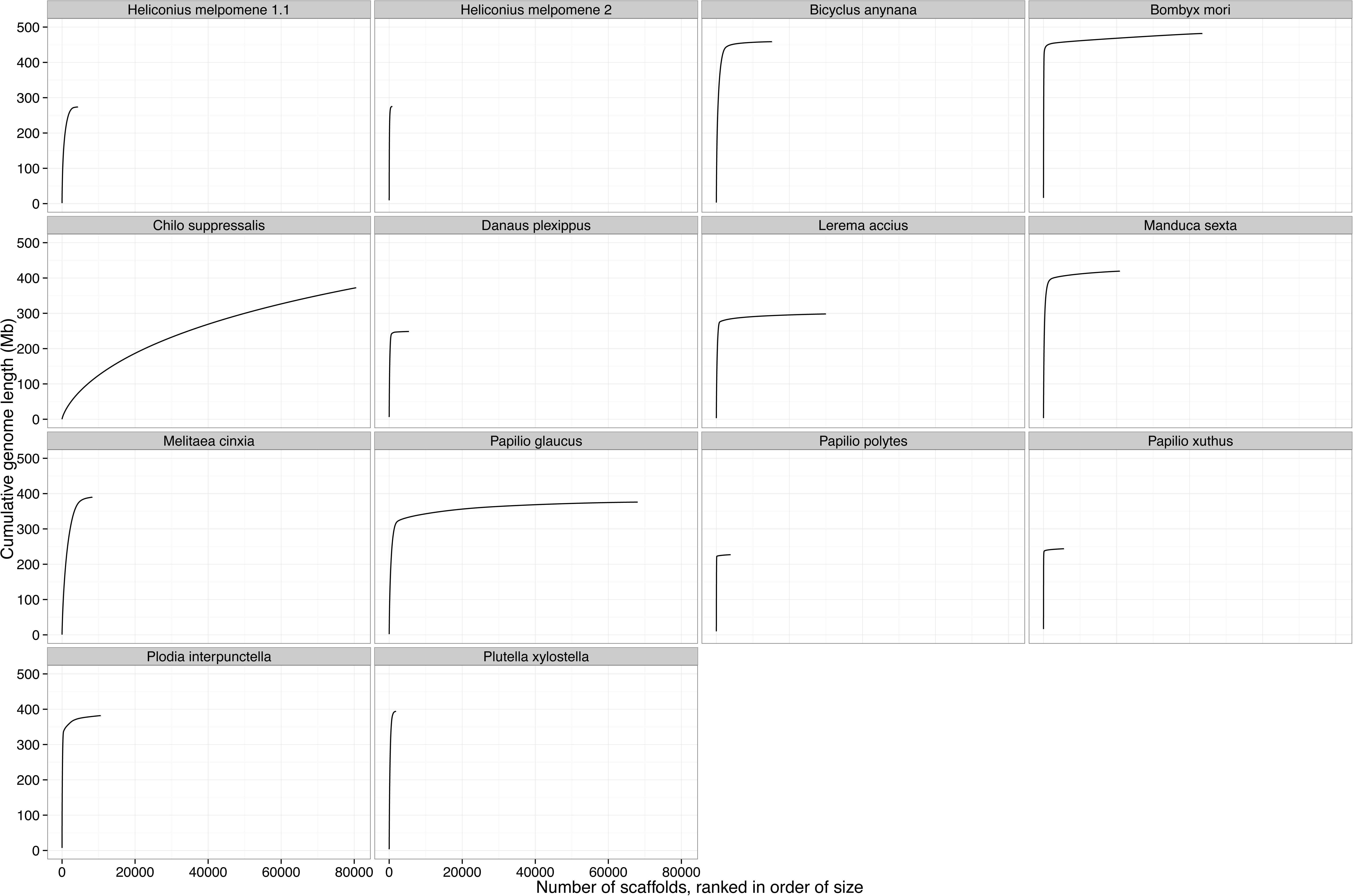
Genome assembly qualities as per Figure 1 for known Lepidopteran genome assemblies. *Bicyclus anynana, Chilo suppressalis, Manduca sexta,* and *Plodia interpunctella* are unpublished draft genomes downloaded from LepBase v1.0 (http://ensembl.lepbase.org) and are likely to change by the time of publication.

**Table S1.** Valid marker types used to build linkage map. A and B are alleles, H is heterozygous for A and B. For A and B calls, the allele may be present in one or two copies. Right columns show number of valid SNPs called for each set of valid parental calls, then grouped by linkage and overall type (Maternal, Paternal, Intercross).

| Marker Type | Linkage | F1 Mother | F1 Father | F2 Males | F2 Females | F0 Grandmother | SNPs |  |  |
| --- | --- | --- | --- | --- | --- | --- | --- | --- | --- |
| Maternal | Autosomal | H | A | A,H | A,H | A | 196,740 | 607,457 | 612,768 |
|  |  |  |  |  |  | H | 410,717 |  |  |
|  | Z-linked | A | B | H | B | B | 5,062 | 5,062 |  |
|  | Pseudo-autosomal | H | A | A | H | H | 126 | 249 |  |
|  | Pseudo-autosomal | H | A | H | A | A | 123 |  |  |
| Paternal | Autosomal | A | H | A,H | A,H | A | 211,171 | 696,187 | 717,758 |
|  |  |  |  |  |  | B | 485,016 |  |  |
|  | Z-linked | A | H | A,H | A,B | B | 15,499 | 21,571 |  |
|  |  |  |  |  |  | A | 6,072 |  |  |
| Intercross | Autosomal | H | H | A,2H,B | A,2H,B | A | 1,157,246 | 1,625,563 | 1,625,563 |
|  |  |  |  |  |  | H | 468,317 |  |  |
| TOTAL | Autosomal |  |  |  |  |  | 2,929,207 |  |  |
|  | Z-linked |  |  |  |  |  | 26,633 |  |  |
|  | Pseudo-autosomal |  |  |  |  |  | 249 |  |  |
|  | ALL |  |  |  |  |  | 2,956,089 |  |  |

**Table S2.**
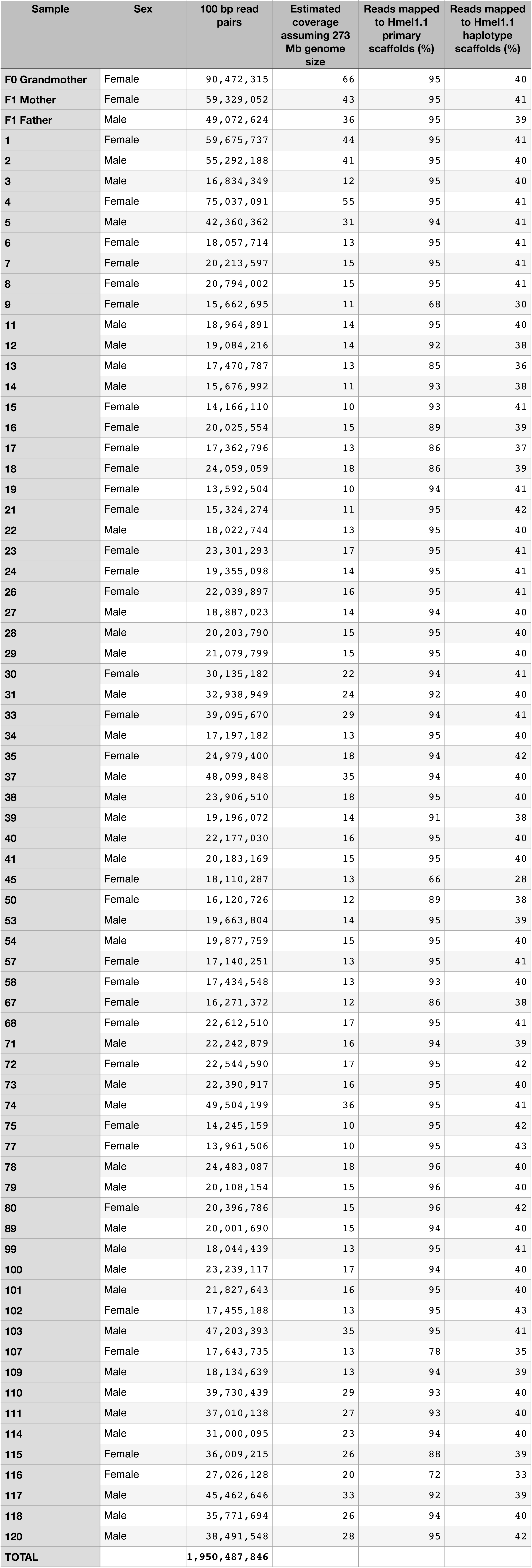
Reads sequenced and mapped for each *H. melpomene* cross individual. Coverage calculated relative to Hmel1.1 genome size of 273 Mb.

**Table S3.** Statistics for PacBio read sets. Combined corrected sets is a merge of ‘Illumina +454 corrected’ and ‘Self corrected’.

| Read set | Reads | Mean length | Maximum length | Total bases | Estimated coverage assuming 273 Mb genome |
| --- | --- | --- | --- | --- | --- |
| Filtered subreads | 1,679,169 | 3,764 | 50,704 | 6,321,685,114 | 23 |
| Illumina+454 corrected | 1,577,076 | 2,644 | 27,868 | 4,170,152,996 | 15 |
| Self corrected | 1,139,019 | 2,947 | 31,243 | 3,357,774,198 | 12 |
| Combined corrected sets | 2,716,095 | 2,772 | 31,243 | 7,527,927,194 | 27 |

**Table S4.**
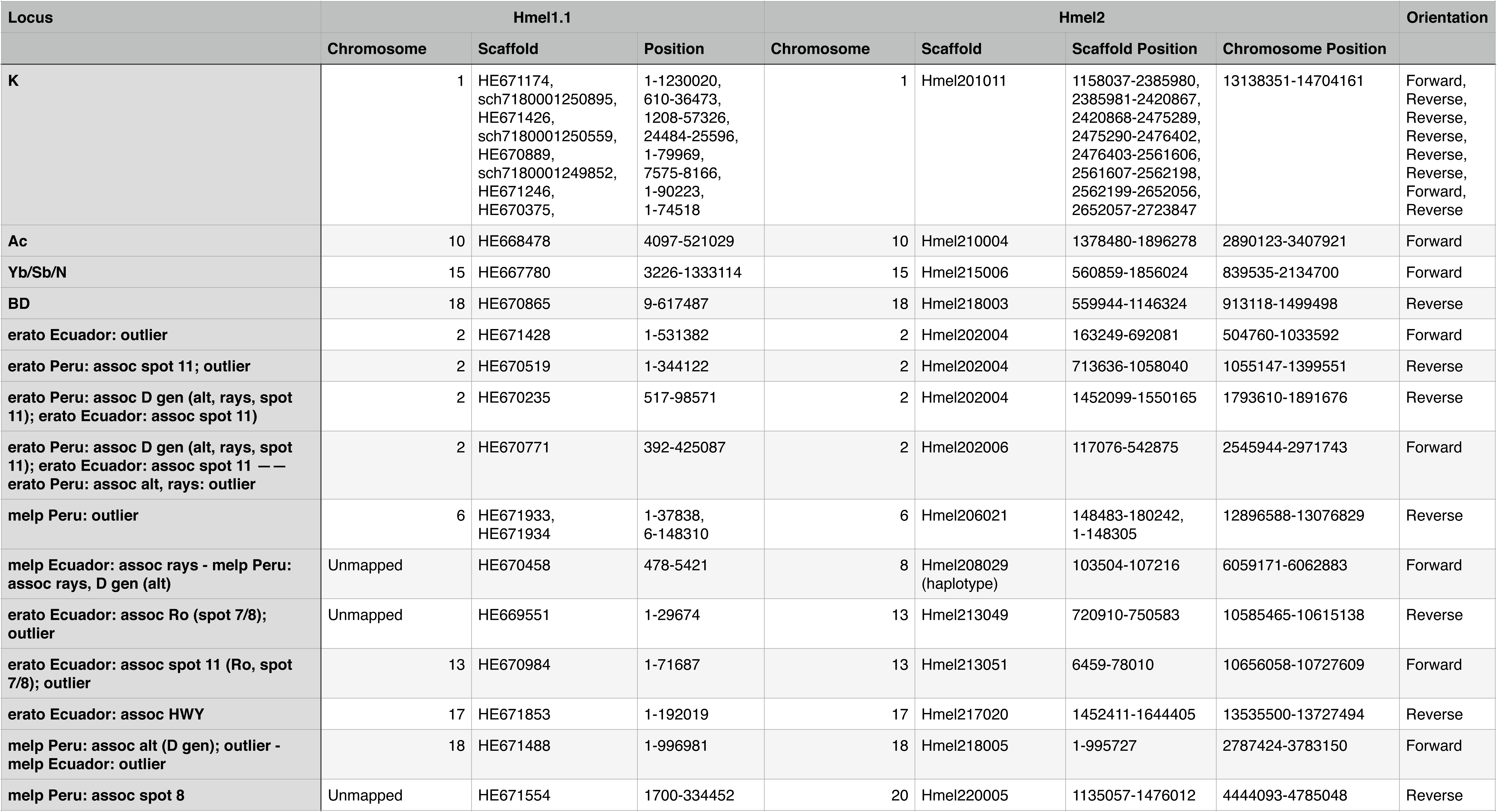
Locations of major adaptive loci from Nadeau *et al.* (2014) in Hmel1.1 and Hmel2. Hmel2 Scaffold Positions refer to the location of the corresponding Hmel1.1 scaffold part in Hmel2, with final orientation of each Hmel1.1 scaffold part given in the Orientation column. Hmel2 Chromosome Positions refer to the location of the entire locus. Hmel1.1 scaffold names begin with ‘HE’ for primary scaffolds and ‘sch’ for haplotype scaffolds.

**Table S5.** Reads sequenced and mapped for the *Eueides isabella* mapping family. All samples were PstI RAD sequenced except for the parents and one offspring, whole genome sequenced for use in a separate project.

| Sample | 151 bp read pairs | Reads mapped to Hmel2 (%) |
| --- | --- | --- |
| MC14-04 (Father) | 84,076,972 | 46 |
| MC14-11 (Mother) | 87,892,825 | 47 |
| MC14-64 | 5,452,089 | 32 |
| MC14-65 | 2,926,927 | 36 |
| MC14-70 | 3,730,478 | 35 |
| MC14-72 | 1,880,271 | 37 |
| MC14-76 | 2,787,303 | 33 |
| MC14-77 | 3,981,681 | 36 |
| MC14-78 | 5,041,043 | 35 |
| MC14-79 | 3,797,695 | 37 |
| MC14-81 | 3,200,779 | 36 |
| MC14-84 | 4,306,783 | 35 |
| MC14-85 | 3,646,291 | 37 |
| MC14-87 | 3,406,938 | 37 |
| MC14-170 | 5,378,843 | 35 |
| MC14-171 | 4,186,544 | 36 |
| MC14-179 | 3,643,677 | 35 |
| MC14-201 | 3,781,326 | 36 |
| MC14-202 | 61,654,236 | 46 |
| MC14-203 | 4,903,067 | 34 |
| MC14-206 | 4,651,268 | 35 |
| MC14-369 | 3,229,381 | 38 |
| MC14-371 | 3,797,574 | 38 |

**Table S6.**
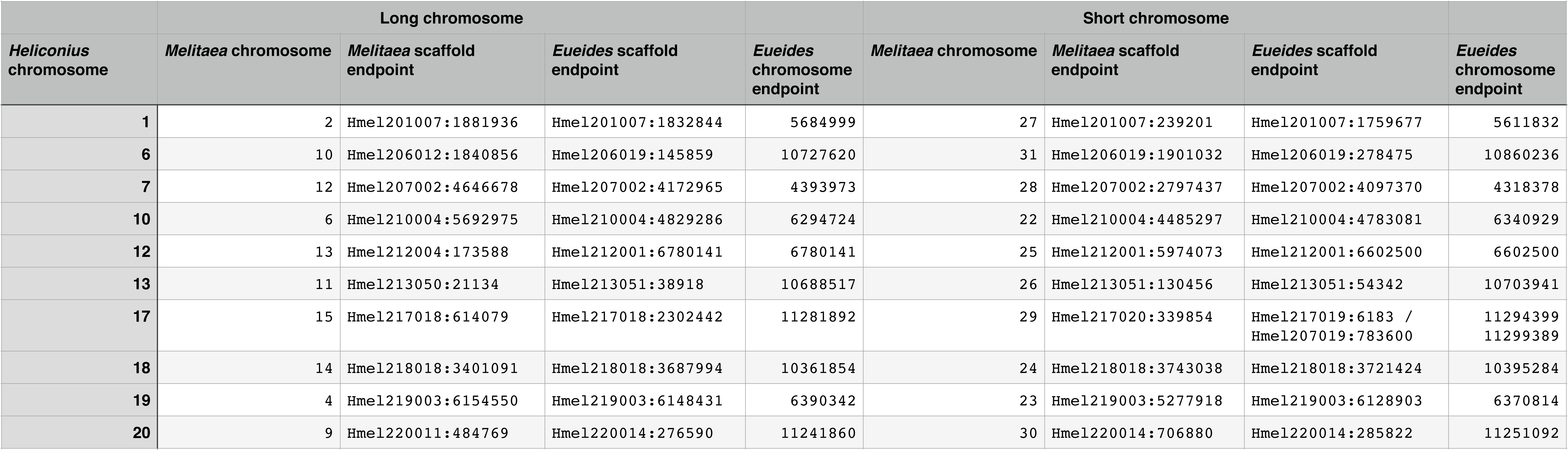
Fusion points for the ten fused *H. melpomene* chromosomes. For each fused chromosome, the two original chromosomes are ordered by length (long on the left, short on the right). Endpoints are positions in the Hmel2 genome. All fusion points fall on single anchored scaffolds except for *H. melpomene* chromosome 17, where the fusion points spans two scaffolds, Hmel217018 and Hmel217019. Hmel217019 is unoriented on the chromosome, so the fusion could end at either end of this scaffold; both possible possible endpoints are given.

| Table S6 |  |  |  |  |  |  |  |  |
| --- | --- | --- | --- | --- | --- | --- | --- | --- |
|  | Long chromosome |  |  |  | Short chromosome |  |  |  |
| *Heliconius* chromosome | *Melitaea* chromosome | *Melitaea* scaffold endpoint | *Eueides* scaffold endpoint | *Eueides* chromosome endpoint | *Melitaea* chromosome | *Melitaea* scaffold endpoint | *Eueides* scaffold endpoint | *Eueides* chromosome endpoint |
| 1 | 2 | Hmel201007:1881936 | Hmel201007:1832844 | 5684999 | 27 | Hmel201007:239201 | Hmel201007:1759677 | 5611832 |
| 6 | 10 | Hmel206012:1840856 | Hmel206019:145859 | 10727620 | 31 | Hmel206019:1901032 | Hmel206019:278475 | 10860236 |
| 7 | 12 | Hmel207002:4646678 | Hmel207002:4172965 | 4393973 | 28 | Hmel207002:2797437 | Hmel207002:4097370 | 4318378 |
| 10 | 6 | Hmel210004:5692975 | Hmel210004:4829286 | 6294724 | 22 | Hmel210004:4485297 | Hmel210004:4783081 | 6340929 |
| 12 | 13 | Hmel212004:173588 | Hmel212001:6780141 | 6780141 | 25 | Hmel212001:5974073 | Hmel212001:6602500 | 6602500 |
| 13 | 11 | Hmel213050:21134 | Hmel213051:38918 | 10688517 | 26 | Hmel213051:130456 | Hmel213051:54342 | 10703941 |
| 17 | 15 | Hmel217018:614079 | Hmel217018:2302442 | 11281892 | 29 | Hmel217020:339854 | Hmel217019:6183 /<br>Hmel207019:783600 | 11294399<br>11299389 |
| 18 | 14 | Hmel218018:3401091 | Hmel218018:3687994 | 10361854 | 24 | Hmel218018:3743038 | Hmel218018:3721424 | 10395284 |
| 19 | 4 | Hmel219003:6154550 | Hmel219003:6148431 | 6390342 | 23 | Hmel219003:5277918 | Hmel219003:6128903 | 6370814 |
| 20 | 9 | Hmel220011:484769 | Hmel220014:276590 | 11241860 | 30 | Hmel220014:706880 | Hmel220014:285822 | 11251092 |

